# Dynamic attention signaling in V4: relation to excitatory/inhibitory cell class and population coupling

**DOI:** 10.1101/2022.08.03.502634

**Authors:** Elizabeth M. Sachse, Adam C. Snyder

## Abstract

Neurons have different roles in attention processing. These roles are determined by a neuron’s firing properties, neurotransmitter expression, and functional connectivity. Neurons in the visual cortical area, V4, are reliably engaged by selective attention but exhibit differences in firing rate and correlated variability. It remains unclear what specific neuronal properties shape these attention effects. We identified neurons as fast-spiking (FS) (putative inhibitory) and regular-spiking (RS) (putative excitatory) and investigated their role in anticipatory attention and how this related to their functional connectivity. V4 neurons exhibited a continuum of time-varying attention effects ranging from “restless-weak” neurons at one extreme to “quiet-strong” neurons at another. We found an interdependence between neural attention effects (e.g., restless-weak or quiet-strong), neuron type (FS, RS), and functional connectivity. In particular, we found neurons with restless-weak attention effects were more likely to be RS and have greater population coupling, compared to neurons with other types of attention effects. Also, quiet-strong neurons were more likely to be FS and these FS neurons exhibited higher spike synchrony. From this we propose that time-varying attention effects in a neuronal population depends on the relative involvement of neurons that drive stimulus processing and those that are engaged by intrinsic population activity. These results add important information to our understanding of visual attention circuits at the cellular level.

## Introduction

Studies of neural activity show diverse attentional effects in sensory cortical regions (Agetsuma *et al*., 2018; Nandy *et al*., 2017; Okun *et al*., 2015; Snyder *et al*., 2016). Specifically, visual area V4 is especially modulated with attention and may function to integrate bottom-up sensory input with top-down cortical modulation (Chelazzi *et al*., 2010; Mehta *et al*., 2000a,b; Snyder *et al*., 2017). V4 is composed of feature and spatial domains that are anatomically and/or functionally connected and extract and integrate sensory information to enable attention (Roe *et al*., 2012). This hypothesis indicates that neuron-type and circuitry of these domains are important. V4 also has layer-specific differences in processes such as orientation tuning (Ringach *et al*., 1997) and attention-related correlation and coherence (Nandy *et al*., 2017). Together, these characteristics make V4 a favorable target to investigate the relationship between neuron-type, local circuit organization, and attention. A central question in neuroscience concerns how diverse effects of attention on neural activity covaries with other neuronal properties, such as whether neurons are excitatory/inhibitory or how those neurons are organized into circuits (Chelazzi *et al*., 2010; Thiele & Bellgrove, 2018) A useful way to answer this question is by identification of neurons based on characteristics of their action potentials. Classification of neurons as FS (putative inhibitory; sometimes also termed “narrow spiking” in the literature) and RS (putative excitatory; sometimes also termed “broad spiking”) by their waveform shapes is an established method in both rodents (Barthó *et al*., 2004; McCormick *et al*., 1985) and macaques (Anderson *et al*., 2013; Nandy *et al*., 2017; Snyder *et al*., 2016). For example, using waveform classification, researchers found that attention modulation acts more strongly on FS neurons (putative interneurons; Mitchell *et al*., 2007; Snyder *et al*., 2016), as compared to RS neurons. In addition, FS neurons thought to express GABA are preferentially targeted by attention signals and induce rapid changes in correlated variability (Snyder *et al*., 2016; Tiesinga *et al*., 2004). Further, optogenetic inactivation of parvalbulmin-expressing interneurons modulates the time-varying attention effects of neuronal ensembles (Agetsuma *et al*., 2018). Taken together, these findings suggest FS and RS neuron types contribute different roles to the circuitry of attention, but the nuances of that division of labor remain poorly understood.

In addition to the neurochemical identities and firing properties of neurons, their functional connectivity (i.e., statistical dependence) is also key to understanding attention processing. For example, different attention modulation of excitatory and inhibitory neurons cannot simply be explained by differences in neural firing rates, instead, it is better explained by changes in correlated variability (Mitchell *et al*., 2007). While much work has gone into identifying the functional connectivity of attention circuits in rodents, it is unclear how this applies to primates. Rhesus macaques have a similar visual system to humans and are capable of performing attention-demanding tasks, but systematic manipulations and circuit identification are more difficult than in rodents due to higher numbers of neurons and limitations of available experimental techniques (Thiele & Bellgrove, 2018; Vinck *et al*., 2013). One way to study functional connectivity in primates is by quantifying neural synchrony, because two neurons that reliably spike around the same point in time are by definition functionally connected. Because inhibitory (FS) neurons are more strongly modulated with attention (Agetsuma *et al*., 2018; Mitchell *et al*., 2007; Snyder *et al*., 2016), one reasonable prediction is that these FS neurons play a larger role in driving attention-mediated neural synchronization compared to excitatory (RS) neurons. To study functional connectivity across groups of neurons, a metric of “population coupling” may be used. Population coupling quantifies the degree to which the activity of one neuron statistically depends on the activity of a broader population of neurons. In an attempt to reconcile the gap in knowledge between rodent and primate attentional circuitry, Okun *et al*. (2015) manipulated neural activity with optogenetics in mice and compared population coupling to that measured by electrophysiology in primates. Those authors demonstrated population coupling is a robust measure that explains patterns of neural activity. They proposed there is an important allocation of labor during stimulus processing among neurons with diverse population coupling ranging from “soloists” (neurons that are not strongly coupled to the population) to “choristers” (neurons that are tightly coupled to population activity). Similar to the findings for FS/RS neurons, “soloists” and “choristers” were found to be influenced to different degrees by internal cognitive factors such as eye-movement planning.

In previous work, we identified FS and RS neurons with distinct attention modulation during a spatial attention task (Snyder *et al*., 2016). FS neurons exhibited greater modulation of firing rates and trial-to-trial variability with attention compared to RS neurons. For the current study, we used a completely data-driven approach to dissect the functional circuitry of attention by investigating the roles of different neuron types in attention over time from the spontaneous to the stimulus-evoked state, and relating those roles to the functional connectivity of individual neurons in the population.

To further parse out the circuitry and neuron-types that generate time-varying attention effects we recorded neural activity in V4 of two rhesus macaques while they performed a spatial attention task. Using a data-driven approach, we identified a spectrum of time-varying attention effects and hypothesized this may be due to the neuron-type composition in a sub-population. The spectrum was characterized by differences in how attention impacted spontaneous versus stimulus-evoked activity. At one end of the spectrum, spontaneous activity was relatively quiet when attended compared to when unattended, but stimulus responses were relatively stronger (we term this the “quiet-strong” pattern); the other end of the spectrum showed the opposite pattern, with relatively high spontaneous activity (which we term “restless”) and weaker stimulus responses (the “restless-weak” pattern).

Using waveform classification, we found quiet-strong neurons were more likely to be FS whereas restless-weak neurons were more likely to be RS. To characterize functional connectivity, we built upon the population coupling approach as by Okun *et al*. (2015). Neurons that tended to be suppressed by attention during stimulus processing were more tightly coupled to the population activity, indicating important differences in functional connectivity based on the role of a neuron in attention. We propose the time-varying attention effects in a given sub-population are generated by the relative presence of FS “soloists” that drive stimulus processing and RS “choristers” that are sensitive to changes in intrinsic population activity. This is an important step towards understanding the circuitry of attention at the cellular level.

## Results

We recorded neural activity in V4 while monkeys performed a visuospatial selective attention task (Snyder *et al*., 2018). Two male rhesus macaques (*Macaca mulatta*; *Monkey P* and *Monkey W*) were trained to detect changes in orientation of Gabor patches in a spatial attention paradigm (Figure 1). One of two stimulus locations was block-cued to be more likely to change (the valid target location). Animals were more accurate at detecting near-threshold targets (Δ3°) at the valid location than at the invalid location, confirming that they selectively attended to the valid location (Monkey P: cue-valid *d*′= 1.76±0.20 [mean ± SEM], cue-invalid *d*′= 0.47±0.20, *t*_23_ = 4.64, *p* = 1.13*·*10^−4^; Monkey W: cue-valid *d*′= 0.46 ± 0.18, cue-invalid *d*′= −0.75 ± 0.18, *t*_22_ = 5.22, *p* = 3.12 *·* 10^−5^). Animals were also faster at detecting targets at the valid location than at the invalid location (Monkey P: reaction time (RT) benefit for cue-valid compared to cue-invalid targets Δ*RT* = −29.2 ± 1.6 ms, *t*_23_ = −17.90, *p* = 5.32 *·* 10^−15^ Monkey W: Δ*RT* = −42.3 ± 4.8 ms, *t*_22_ = −8.79, *p* = 1.19 *·* 10^−8^).

**Figure 1:**
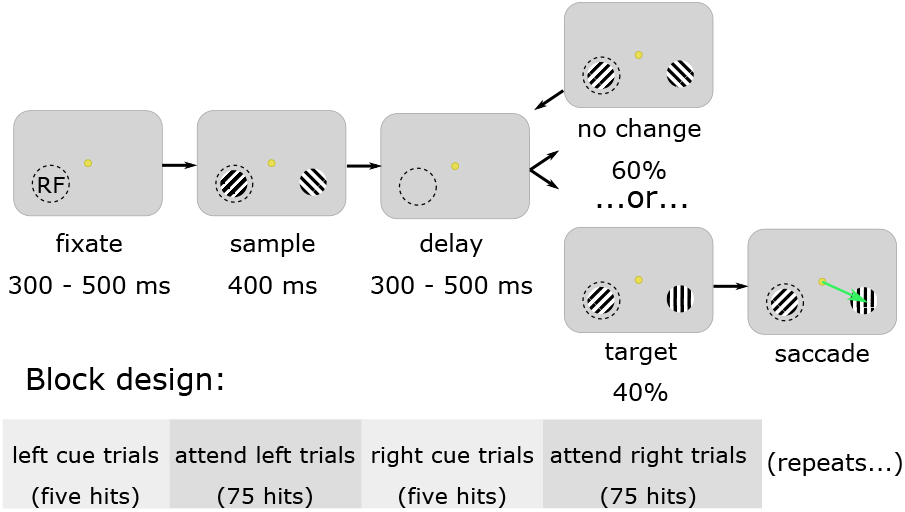
Task design. Subjects initiated each trial by fixating their gaze on a central yellow dot. After a fixation interval, Gabor stimuli were repeatedly flashed for 400ms, separated by 300–500ms inter-stimulus intervals. One of the stimuli covered the aggregate RF area of the recorded V4 population. The subjects’ task was to detect a change in orientation of one of the two stimuli (the target) and make a saccade to the stimulus that changed. Each stimulus flash had a fixed chance of containing a target (30% for monkey P, 40% for monkey W). One location had a 90% chance of containing the eventual target (the valid location), and we alternated the side of high-target probability after 80 correct detections (hits). For the initial trials in each block only one stimulus was presented at the valid location (cue trials) until the subject made 5 correct detections, after which bilateral stimuli were presented for the remainder of the block. The dashed circle represents the receptive field (RF) and was not actually present in the display.

Separate analyses of portions of these data were previously reported (Snyder *et al*., 2018; Cowley *et al*., 2020; Snyder *et al*., 2021; Umakantha *et al*., 2021; Johnston *et al*., 2021). Here, we tested the degree to which attention dynamics were related to sub-populations of neurons that differ in neuron-type and functional connectivity.

### Relationship between neuron-type and time-varying attention effects

The first question we sought to answer was: do different time-varying attention effects relate to functional properties of the neurons in the population? We compared the attention signals of single-units (Example units shown in Figure 2). In the peristimulus spike-time histograms (PSTHs) a distinct pattern emerged. The firing rates of some neurons were relatively suppressed by attention following stimulus onset (Figure 2c, 2d) whereas the firing rates of other neurons were enhanced by attention following stimulus onset (Figures 2a, 2b). Throughout this report, we refer to the time-course of attention modulations such as these (e.g., Figure 2b,d) as “time-varying attention effects”.

**Figure 2:**
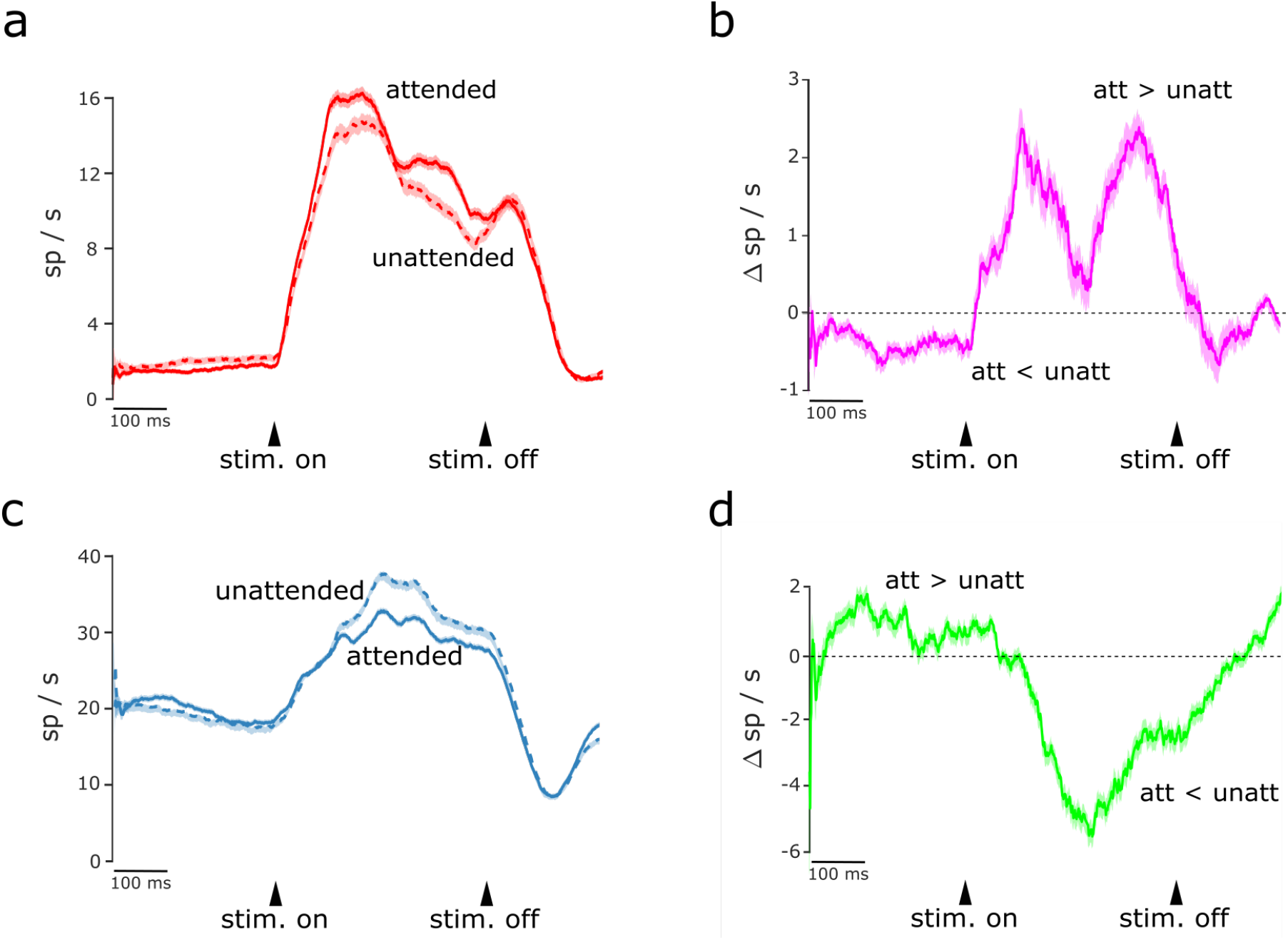
PSTH and attention modulation for two example neurons. **(a)** A PSTH from an example neuron whose pre-stimulus activity was higher for the unattended condition compared to the attended condition, and whose post-stimulus activity was conversely higher for the attended condition (units are in spikes per second). The SNR of this example neuron = 4.22. Shading indicates the standard error of the mean (SEM). **(b)** Attention modulation (attended responses minus unattended response) for the example neuron in (a). Units are the difference in spikes per second of the attended versus unattended condition. Shading indicates SEM. **(c)** A PSTH from an example neuron showing an opposite pattern compared to (a). Pre-stimulus activity was higher for the attended condition than the unattended condition, and post-stimulus activity was conversely higher for the unattended condition. The SNR of this example neuron = 2.78. **(d)** Attention modulation for the example neuron in (c).

To test if the observed patterns of time-varying attention effects were isolated to certain neuron-types we next classified neurons as FS or RS. To do this, we used an automated clustering algorithm based on waveform shapes (as in Snyder *et al*., 2016). We trained our algorithm on the best-isolated neurons (those with signal-to-noise ratios [SNRs] in the top quartile of the sample; see Methods), and then applied the trained algorithm on all neurons, regardless of SNR. We observed that neurons with low SNR tended to be classified as FS at a greater rate than neurons with high SNR (> 3; Figure 3), which we reasoned was likely a limitation of the automated clustering method rather than a biological finding. For this reason, we performed subsequent analyses on a restricted sample of neurons with SNRs greater than 3 (unless otherwise indicated). For completeness and comparison, we also performed our analyses on the full sample of neurons (i.e., SNRs > 0). As we shall describe, the patterns of results were generally similar for our full sample of neurons as for the restricted sample of high-SNR neurons. The majority of neurons in each neuron-type group had significant differences in firing rate between attention conditions (FS, SNR unrestricted: 78% ±1.7% (95% CI); FS, SNR restricted: 83% ±3.2%; RS, SNR unrestricted: 80% ±2.2%; RS, SNR restricted: 80% ±3.2%; Figure 3). This demonstrates that both FS and RS neurons were modulated with attention. Our second question was: were there distinct classes of time-varying attention effects or were they continuously distributed? Identifying types of neurons involved in visual and attentional processing is a common research interest (Agetsuma *et al*., 2018; Mitchell *et al*., 2007; Nandy *et al*., 2017; Okun *et al*., 2015; Snyder *et al*., 2016). Since we found variation in attention dynamics, a natural question is, did the diversity of dynamics map onto classes of neurons already identified? If time-varying attention effects were related to specific neuron classes we would predict to find clusters of time-varying attention effects. Alternatively, if time-varying attention effects were not clustered, this would suggest a subtler relationship between time-varying attention effects and other neural properties, or no relationship at all.

**Figure 3:**
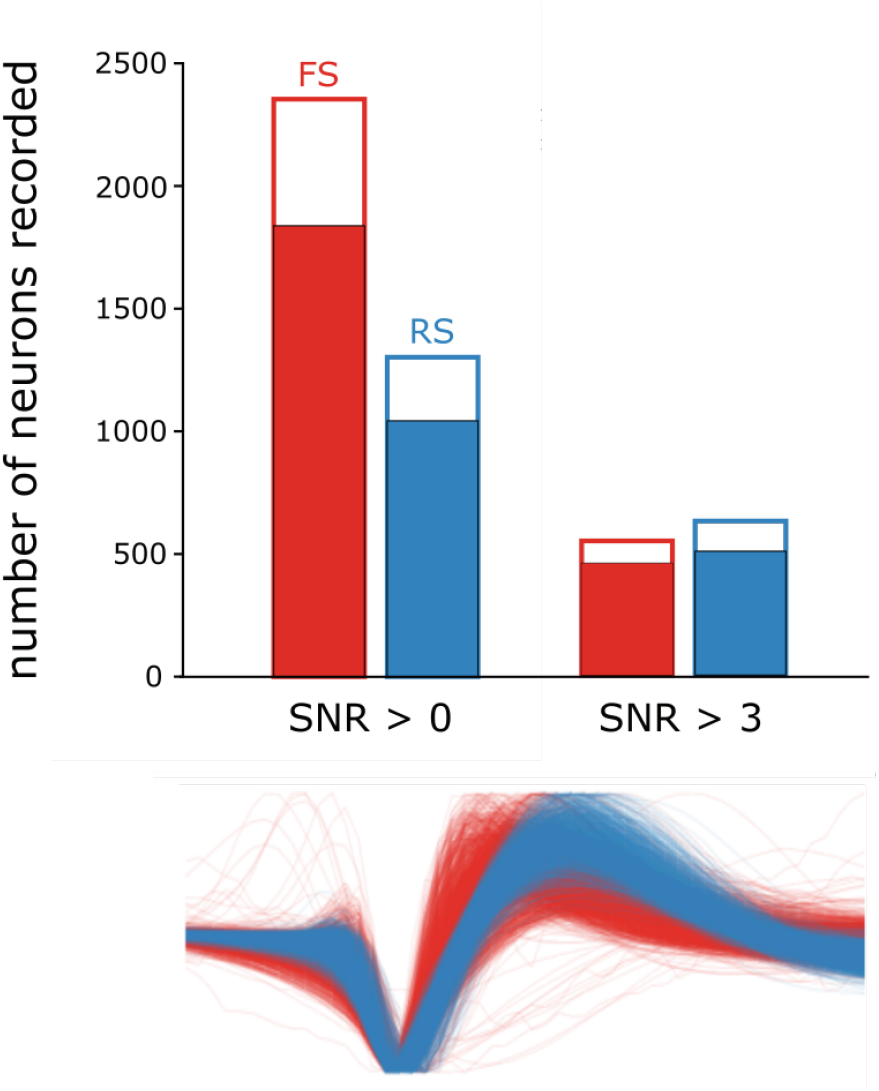
The total number of classified neurons (filled portion indicates neurons with significant attention effects). Below, amplitude-normalized spike waveforms for FS (red) and RS (blue) neurons.

To test for clustering, we used an unsupervised, data-driven approach to reveal any potential structure in the distribution of time-varying attention effects. We first quantified the time-varying attention effects of each neuron by an “attention index” ([attended-unattended]/unattended). We next measured Pearson distances between time-varying attention effects for all pairs of neurons and applied multi-dimensional scaling (MDS) as a dimensionality reduction method to approximate those relative distances in two dimensions (Figure 4a). The result of these steps is a two-dimensional embedding of our sample of neurons where the distance between two neurons is inversely related to the similarity between the time-varying attention effects of those two neurons.

**Figure 4:**
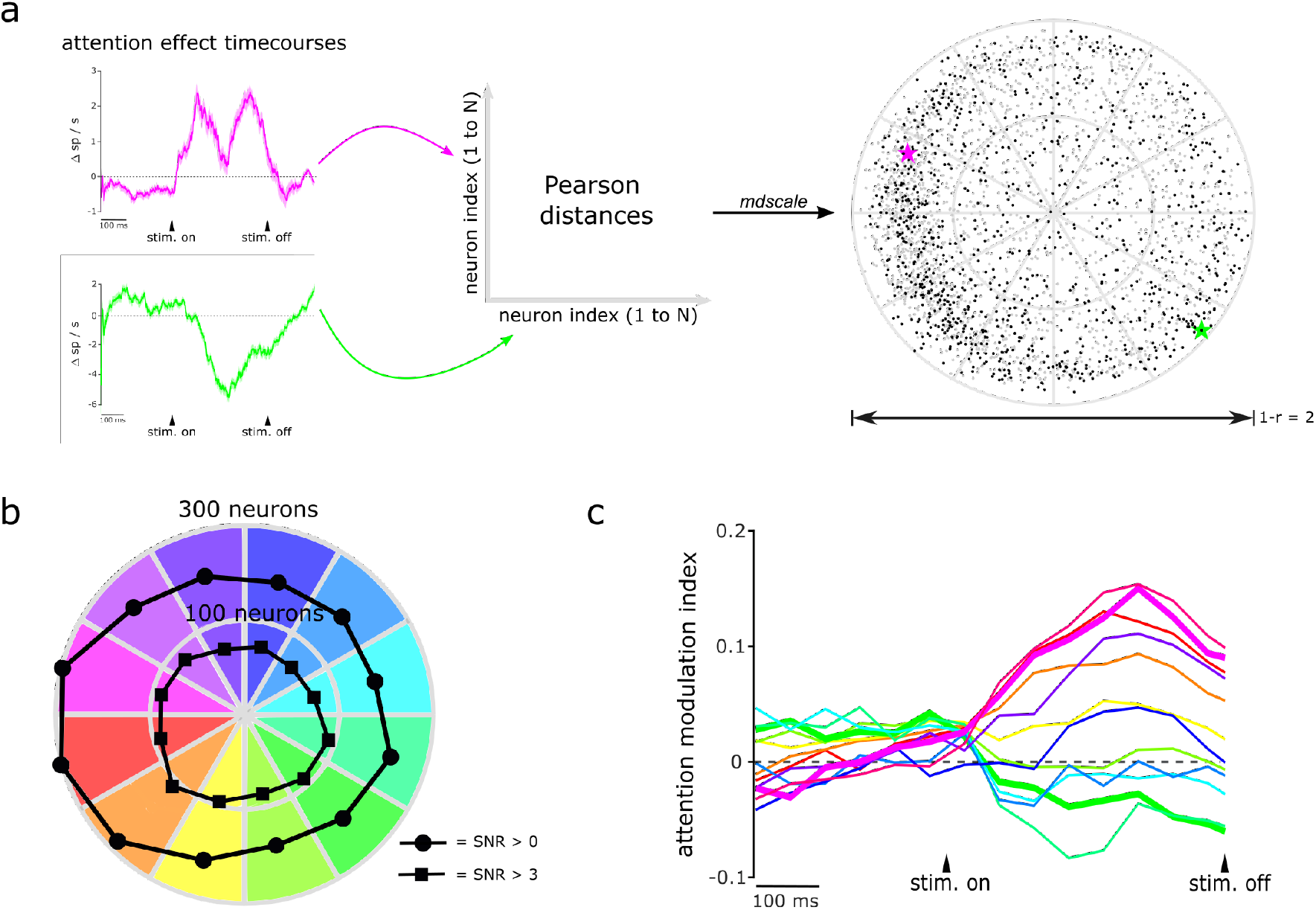
Multi-dimensional scaling (MDS) of attention effect time-courses for neurons with significant attention effects. **(a)** Method to perform MDS and the resulting distribution. A Pearson distance (1 − *ρ*, where *ρ* is the Pearson correlation) was calculated from the attention effect timecourses of each pair of neurons. The two example neurons shown in figure 2 have a Pearson distance of 1.39, indicating highly dissimilar attention effect timecourses. Then, MDS was performed on all Pearson distances to embed the neurons in a two-dimensional space while minimizing distortion of their pairwise Pearson distances, resulting in the distribution shown on the right. Each point represents a neuron; black points indicate the neuron has a SNR > 3, while gray points indicate a SNR > 0. The two example neurons location in MDS space are indicated by stars. The distance between each pair of points is approximately proportional to the Pearson distance between attention effect time-courses for the corresponding neurons. That is, nearby points have similar attention effect time-courses. Since no clustering of points was found (see Methods), we grouped neurons based on arbitrary spatial sectors for further analysis. This would be analogous to e.g. binning up neurons based on a one-dimensional axis such as “attention index”, but generalized to a higher-dimensional space. **(b)** Total number of neurons in each MDS sector (radius of polar plot). **(c)** Attention modulation index [(attended-unattended)/unattended] for neurons with similar attention effects. Color corresponds to MDS sector in (b). Bold traces represent the two ends of the spectrum of attention effects: pink = quiet-strong and green = restless-weak.

We next asked whether there were distinct “clusters” of time-varying attention effects in the sample, or rather if time-varying attention effects was continuously distributed as a spectrum. To test for clustering, we used the k-means algorithm for various potential numbers of clusters (*k*) and quantified the result using Bayes’ Information Criterion (BIC). We found that the model with *k* = 1 (that is, just a single cluster) performed best, which indicated that time-varying attention effects were continuously distributed as a spectrum, rather than organized into distinct categories (we also tested for clustering in embedding dimensionalities of three, four and five dimensions, and found the same result).

However, to test for subtler relationships between time-varying attention effects and other neuronal properties, we arbitrarily broke up the MDS result into 30° sectors (the qualitative pattern of results and our interpretations were unaffected by moderate variation around this parcellation scheme). That is, we grouped neurons into twelve groups (one for each sector of the MDS space) based on the similarity of their time-varying attention effects (Figure 4). This is analogous to the common practice of dividing up a sample of neurons into “bins” based on some continuously-valued property, and then measuring other properties of the neurons in each bin. Because our MDS result had circular structure with most of the density near the edge, sectors, which are the angular analog to bins, were most reasonable. Examining the average time-varying attention effects in each sector, we saw that at one end of the spectrum, neurons were generally quieter during spontaneous activity when attended, compared to when unattended, but then had stronger stimulus responses when attended, compared to when unattended (“quiet-strong” neurons; Figure 4c pink); at the other end of the spectrum, neurons were generally more active with attention during spontaneous activity, but then had weaker stimulus responses (“restless-weak” neurons; Figure 4c green). Other sectors showed time-varying attention effects that were intermediate between those extremes. While the salient feature of the spectrum of time-varying attention effects turned out to be well-described by differences in attention modulations during spontaneous, as compared to stimulus-evoked activity, it is worth emphasizing that our unsupervised and data-driven approach had the potential to find more complex temporal structure in the data with minimal *a priori* assumptions.

We next tested whether the relative proportion of FS neurons differed across sectors, which would suggest a relationship between time-varying attention effects and neuron type. We calculated the relative odds that a neuron randomly picked from each sector would be FS compared to the odds of picking a FS neuron from the whole sample (i.e., the “odds ratio” that a neuron in each sector is FS). We found that neurons in a sector containing quiet-strong neurons were relatively more likely to be FS (Figure 5, p = 0.0072 for SNR unrestricted, bootstrap test), and that neurons in a sector containing restless-weak neurons were relatively more likely to be RS (Figure 5, p = 0.0004, SNR unrestricted; p < 0.0001, SNR restricted). This suggests there is a relationship between time-varying attention effects and FS/RS neuron types in V4. However, because time-varying attention effects were not co-clustered with neuron types, this relationship would be best described as a subtle trend, rather than a categorical difference.

**Figure 5:**
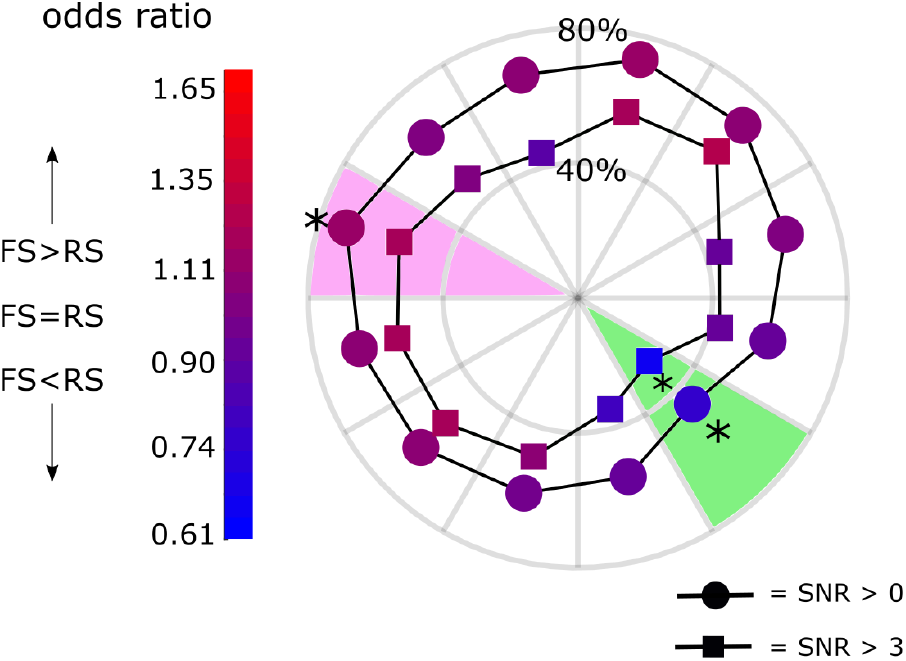
Relationship between neuron type and time-varying attention effects. Radius indicates proportion of FS neurons in each sector. Color indicates odds ratio of a neuron being FS in each sector (values greater than 1 mean FS neurons are over-represented relative to the sample as a whole). Circular markers indicate all neurons with SNR values > 0 were included in the analysis. Square markers indicate only neurons with an SNR > 3 were included. In the quiet-strong sector (pink), for unrestricted SNR values the risk of a neuron being FS was significant: p = 0.0072. In the restless-weak sector (green) neurons were significantly less likely to be FS: p = 0.0004 for unrestricted SNR values and p< 0.0001 for high SNR only neurons (bootstrap test, significance criterion: *α* < 0.1, Šidák-corrected for number of sectors tested.)

### Relationship between population coupling and time-varying attention effects

Next we tested the relationship between time-varying attention effects and pairwise spike synchrony. We predicted that changes in synchrony of specific neuron-types and in specific parts of the attention circuitry could act to enhance attention signals from higher-order areas such as the prefrontal cortex (PFC), and help regulate the behavioral response. In both animals, we found time and neuron-type-dependent effects on spike synchrony. Synchrony was significantly greater prestimulus compared to poststimulus in both animals and across neuron pair types (Figure 6 and Table 1, ANOVA on pooled excess coincidences per spike for each animal, testing main effects and interactions of attention effect, time, and pair type). Average synchrony between FS-FS pairs was higher than RS-RS and mixed pairs for both animals (Figure 6, Tukey multiple comparisons test). This is interesting because FS neurons are typically interneurons that participate in local circuitry to modulate population responses (Agetsuma *et al*., 2018). We found no distinct differences in synchrony between neuron pair types during stimulus processing in either attention condition. However, for Monkey P only, there was a significant interaction effect of time and attention condition on synchrony. Prestimulus, attended neurons were more synchronous than unattended neurons (Figure 6, monkey P). Further, the scale of spike synchrony was slightly different between animals. Regardless of these subtle differences, the pattern of results is the same in both animals: synchrony was greater prestimulus than poststimulus, and greatest in FS neuron pairs. These synchrony results did not capture nor explain features of the time-varying attention effects we observed (Figure 4c). From this we hypothesized synchrony may work through a broader network rather than pair-wise interactions based on a specific neuron-type or attention condition. Therefore, we tested a more global measure of functional connectivity: “population coupling”.

**Table 1:**
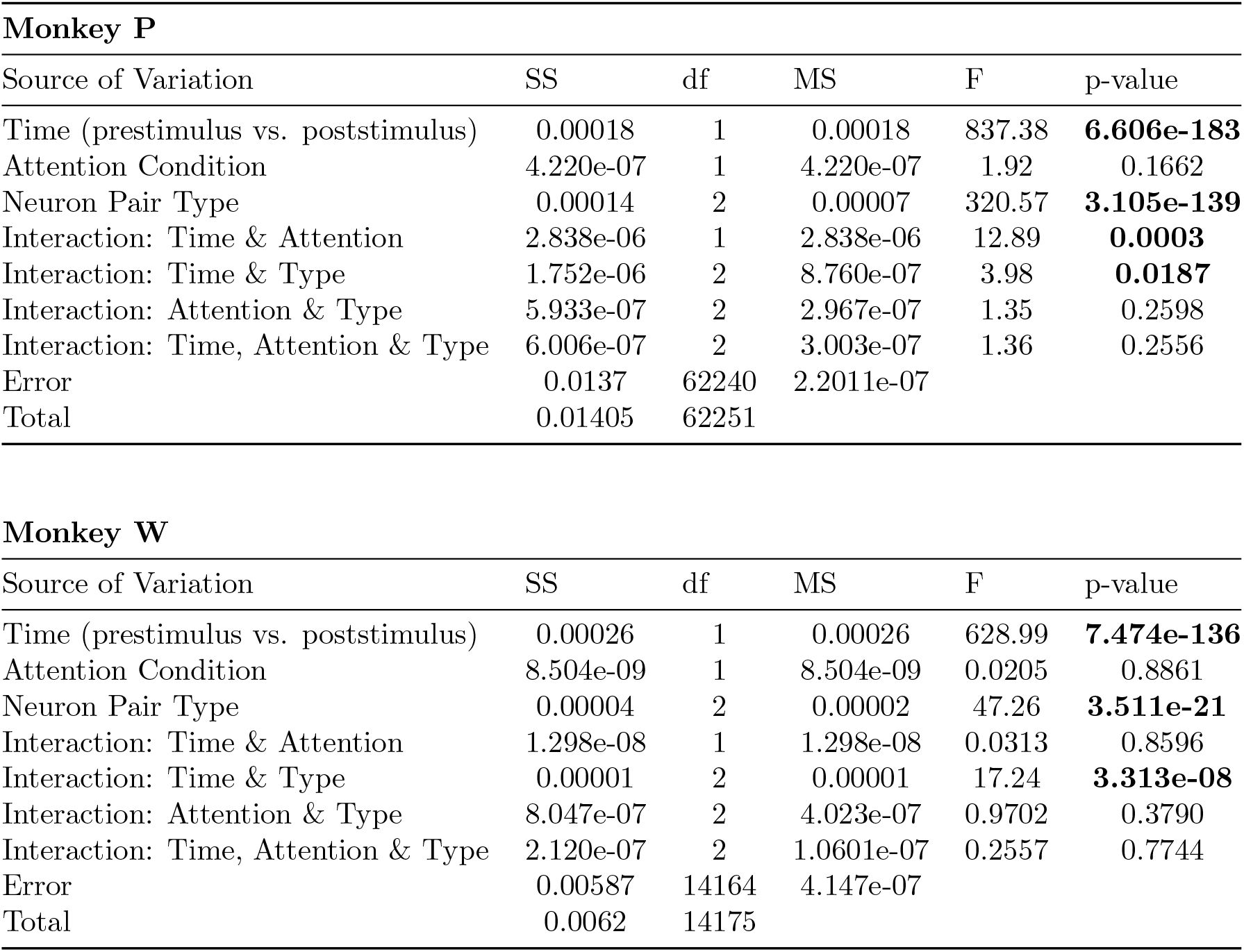
Results from an N-way ANOVA analysis of excess coincidences per spike for each animal. For results from post-hoc multiple comparisons see Figure 6

**Figure 6:**
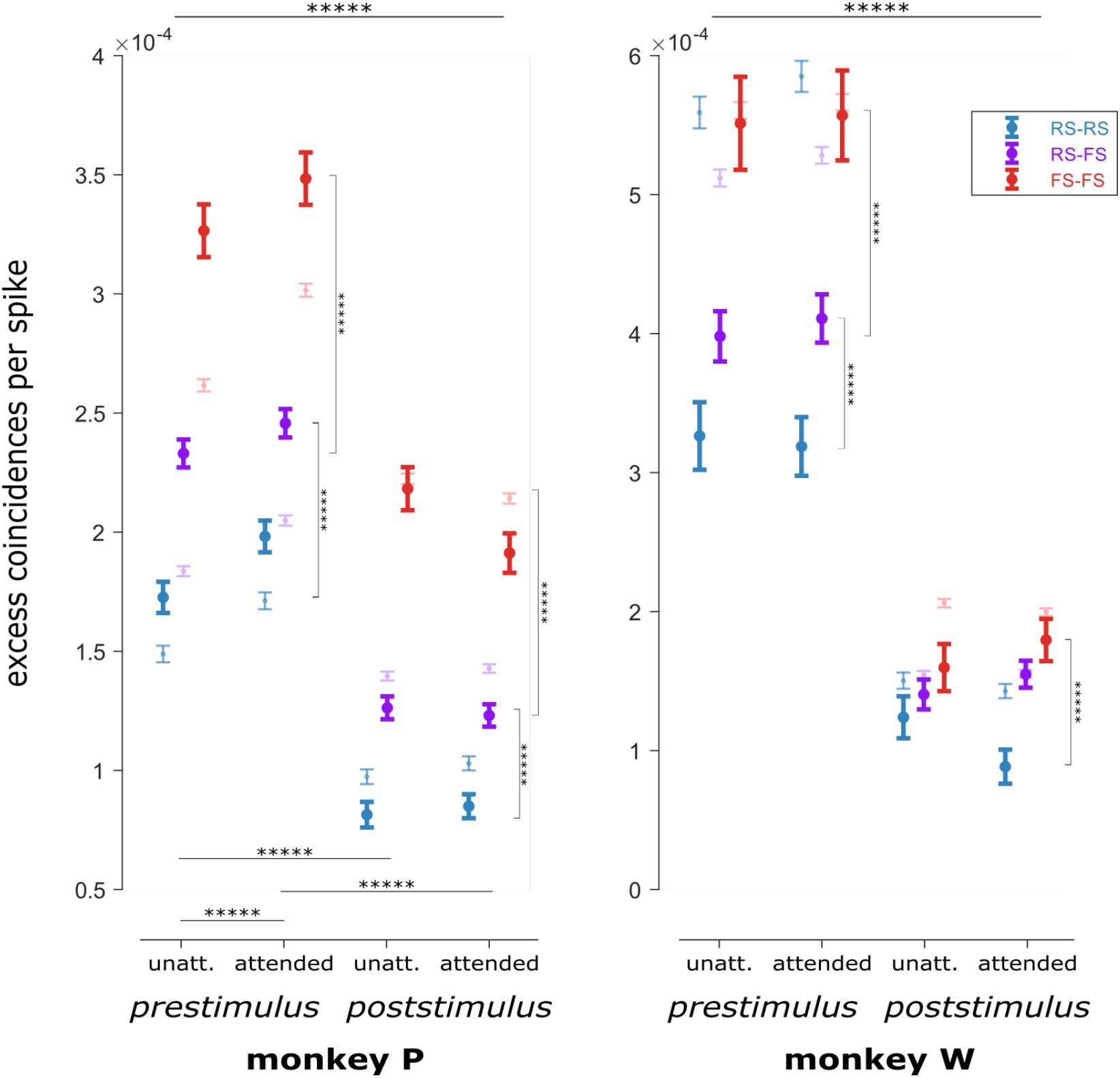
Excess coincidences per spike for all neuron pair types. Faded data points represent synchrony for all neurons while the regular data points represent synchrony for neurons with a SNR > 3. Reported here are statistics for SNR restricted neurons only: Prestimulus and poststimulus synchrony were significantly different for both animals (*p* < 0.0001) as well as synchrony by pair type (*p* < 0.0001). Synchrony was not significantly different between attended and unattended conditions. In both animals, there was a significant interaction between time and neuron type (Monkey P: *p* = 0.0187, Monkey W: *p* < 0.0001). Monkey P also exhibited a significant interaction between time and attention (*p* = 0.0003). For full results of the N-way ANOVA analysis see Table 1. **Monkey P:** Prestimulus and poststimulus, FS-FS pairs were significantly more synchronous than RS-RS pairs and FS-RS pairs (Tukey multiple comparisons test, both *p* values < 1 · 10^−20^). FS-RS pairs were significantly more synchronous than RS-RS pairs prestimulus (*p* = 1.92 · 10^−18^) and poststimulus (*p* = 1.12 · 10^−10^). In addition, prestimulus synchrony was significantly higher than poststimulus synchrony for both unattended and attended neurons (both *p* values < 1 · 10^−20^). Prestimulus, attended neurons were more synchronous than unattended neurons (*p* < 1 · 10^−20^). **Monkey W:** Prestimulus, FS-FS neuron pairs were significantly more synchronous than FS-RS pairs (*p* = 7.79 · 10^−15^) and RS-RS pairs (*p* < 1 · 10^−20^). Poststimulus, FS-FS pairs were more synchronous than RS-RS pairs (*p* = 0.0392) but were not significantly different from FS-RS pairs. Error bars indicate ±1 SEM.

While synchrony is a pair-wise measure between neurons, population coupling is a property of individual neurons. Population coupling describes how the activity of a neuron is statistically dependent with the overall population activity. This measure is sensitive to different attributes of functional connectivity than the conventional analysis of neural synchrony used above, since it takes into account a neuron’s individual activity compared to all other neurons in the sample. One way to quantify population coupling is through centrality measures, i.e., how “centrally” located one neuron is in the functional population space. Here, neurons that have higher synchrony relative to other neurons in the population will be more “central” in the network. To calculate average closeness centrality of the neurons in each MDS sector, we first calculated synchrony (excess coincidences per spike) across all neuron pairs (Figure 7a). From this, we created a graph representation for each sample of neurons (i.e., each recording session), where each neuron is represented as a node and the distance between each pair of nodes is the inverse of the synchrony for that pair (example graph shown in Figure 7b). Closeness centrality can then be calculated for each neuron in this graph representation as the average minimum distance to all other neurons (note that because synchrony as a distance metric does not obey the triangle inequality, the minimum distance between two nodes may be indirect, via a third node). We found that restless-weak neurons were more tightly coupled to the population activity (Figure 7c). In both animals, the sub-population of neurons in the restless-weak sector had significantly higher closeness centrality indicating tighter coupling to the population rate (Figure 7c, permutation test). This sector also had significantly more RS neurons (Figure 5) whose firing rates were enhanced before stimulus onset (Figure 4c, bold green line). Although not statistically significant, the quiet-strong sector, which had significantly more FS neurons (Figure 5) that were suppressed prior to stimulus onset (Figure 4c, bold pink line) had relatively low centrality and therefore lower population coupling. Taken together, higher population coupling for neurons in the restless-weak sector may be associated with “choristers” that are more sensitive to non-sensory information such as top-down attention signals (Okun *et al*., 2015). In contrast, neurons in the quiet-strong sector may have more “soloists’ that process stimulus-related information. This demonstrates a relationship between neuron-type, local connectivity and attention dynamics. We further separated out population coupling in each MDS sector by neuron type. We hypothesized that RS neurons would be more central than FS neurons, especially in the restless-weak sector. Surprisingly, in both monkeys, the restless-weak sector contained FS and RS neurons with high closeness centrality (Figure 7d). In contrast, in the quiet-strong sector, FS and RS neurons similarly exhibited low levels of centrality. The fact that both FS and RS neurons contribute to the high centrality in the restless-weak sector may indicate that there are specialized circuits in V4 composed of both excitatory neurons and local inhibitory neurons that are tightly coupled to the population activity.

**Figure 7:**
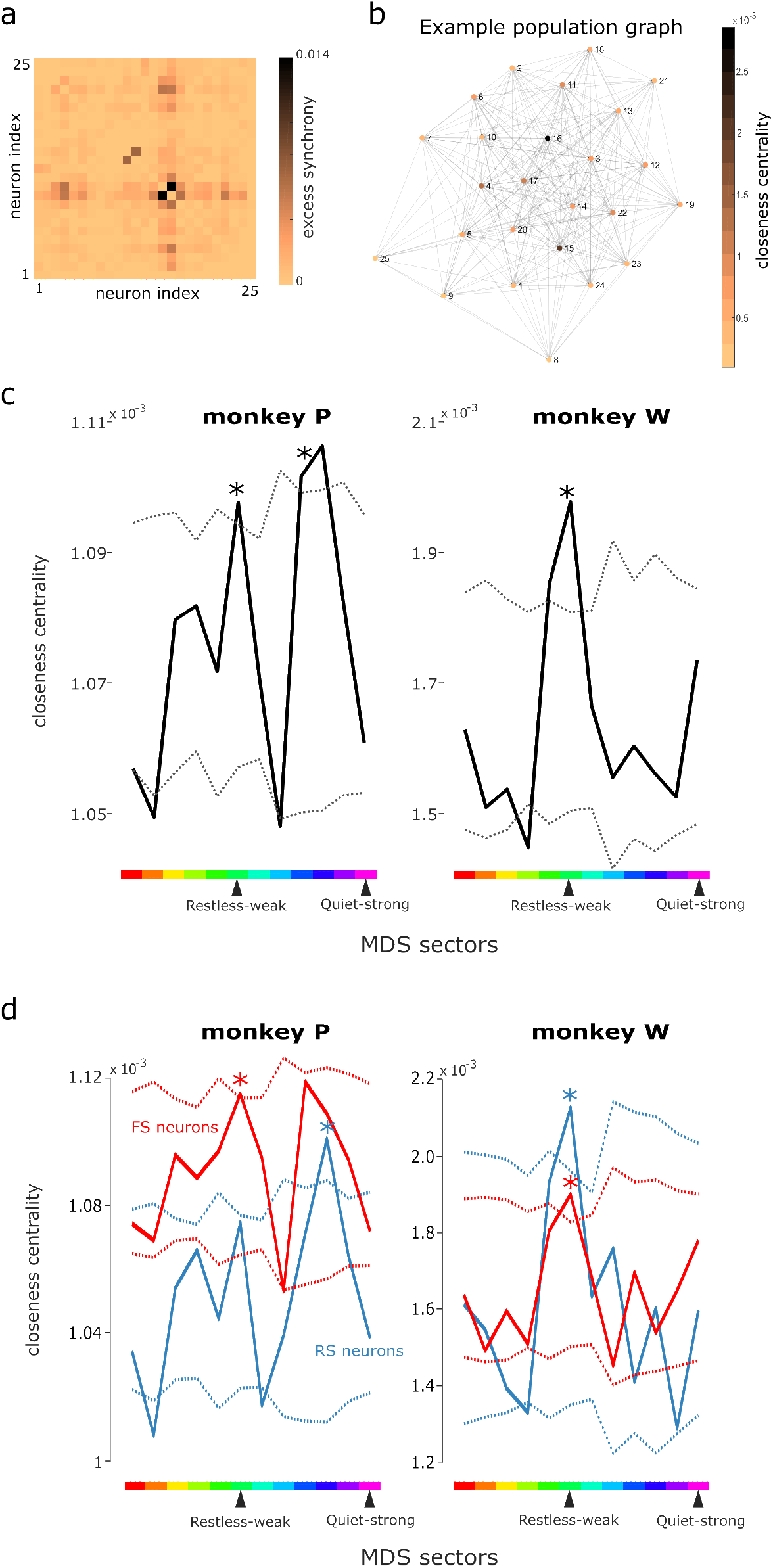
Population coupling **(a)** Excess synchrony between all neurons from an example session. **(b)** Excess synchrony was converted to distance to make a centrality network from the example session in ‘a’. Each node is a single neuron. Neurons that are more synchronous with each other are more centrally located in the network. **(c)** Population coupling over all sessions. Colors along abcissa correspond to MDS bins as colored in Figure 4b. In both monkeys, the sector corresponding to restless-weak timevarying attention effects (green bin on abcissa), had significantly higher closeness centrality than expected by chance. Dashed lines indicate 2.5 and 97.5 percentiles of bootstrap null distribution; values outside this range are significantly more extreme than expected by chance if there was no relationship between sector and centrality. **(d)** Population coupling over all session separated by neuron type. For Monkey P, the sector corresponding to restless-weak time-varying attention effects (green bin on abcissa) contained FS (red line) with significantly higher closeness centrality than expected by chance with RS (blue line) neurons approaching significance. For Monkey W, the green sector corresponding to restless-weak neurons contained both FS and RS neurons with significantly higher closeness centrality. Dashed lines indicate 2.5 and 97.5 percentiles of bootstrap null distribution.

## 1 Discussion

We sought to improve our understanding of the neural circuitry of attention in V4 by testing the association of time-varying attention effects to measures of neuron-type (FS or RS) and population coupling. First, we found a spectrum of time-varying attention effects in V4, with neurons at one end of the spectrum showing suppressed spontaneous activity and enhanced stimulus responses (quiet-strong modulations), and neurons at the other end of the spectrum showing the opposite pattern (restless-weak). Second, although again not a categorical distinction, we found that neurons with quiet-strong modulations had a greater odds ratio of being FS; whereas neurons with restless-weak modulations had a greater odds ratio of being RS. Third, neurons with restless-weak modulations had greater population coupling (as measured by closeness centrality of synchrony) than neurons with other types of attention modulations. Taken together, these findings suggest that time-varying attention effects relate to neurons’ circuit properties, such as whether they are inhibitory vs. excitatory, or weakly vs. strongly coupled to the rest of the population.

During endogenous attention, V4 dynamics are broadly regulated by both endogenous top-down signals from other cortical areas and exogenous sensory inputs via visual circuitry (Roe *et al*., 2012; Pasupathy *et al*., 2020). Based on this, we would expect a variety of neuron sensitivities to endogenous cortical inputs or specific features of the visual stimulus rather than homogeneity. Further, these sub-populations will likely be differently modulated with attention depending on the context of the experimental task, the type of visual stimulus (i.e. Gabors, oriented gratings, moving dots), and the neuron types comprising that sub-population. Also, sensory cortical responses to visual stimuli unfold over time. Although most of the available sensory information in a visual stimulus is contained by the earliest-occurring spikes in response to that stimulus (Osborne *et al*., 2004), feature selectivity in visual cortical areas continues to develop with time (Sani *et al*., 2017).

This dynamic sensory encoding has been observed in multiple sensory cortical regions and for different features of the stimulus (V1: orientation selectivity (Ringach *et al*., 1997) and contour integration (Chen *et al*., 2014), V4: texture selectivity (Kim *et al*., 2019) and object occlusion (Fyall *et al*., 2017), and MT: pattern motion sensitivity (Smith *et al*., 2005)). In sum, complex local circuitry is required to engage the correct combinations of neurons at the appropriate time and extract relevant sensory information to generate dynamics of feature selectivity. The current findings clarify the circuit mechanisms underlying this process.

A recurring observation in attention studies is that FS neurons seem to be preferentially engaged by attention (Agetsuma *et al*., 2018; Anderson *et al*., 2013; Barthó *et al*., 2004; Hofer *et al*., 2011; Mitchell *et al*., 2007; Snyder *et al*., 2016; Shin & Moore, 2019; Tiesinga *et al*., 2004; Vinck *et al*., 2013). The current results add to this growing literature suggesting a specialized role for FS neurons in attention by demonstrating that neurons with quiet-strong time-varying attention effects are relatively more likely to be FS compared to neurons with other patterns of time-varying attention effects. Because FS neurons have been previously implicated as inhibitory (Barthó *et al*., 2004; Mitchell *et al*., 2007; McCormick *et al*., 1985), this deepens our understanding of the functional role for inhibition in attention processes. However, it is important to note that characterization of waveform shape only putatively identifies FS and RS neurons as inhibitory or excitatory, respectively. Further, within these classes there are many sub-types of inhibitory and excitatory neurons with different functional circuitry and response properties (Agetsuma *et al*., 2018; Pritchett *et al*., 2015; Shin & Moore, 2019). In our sample of neurons, we have a high proportion of FS neurons compared to other studies of V4 (Mitchell *et al*., 2007; Nandy *et al*., 2017; Hofer *et al*., 2011; Snyder *et al*., 2016; Vinck *et al*., 2013). This may be because our chronic array method recorded relatively many neural units with low SNR (since electrode position could not be fine-tuned to improve isolation), which we found were typically classified as FS. To account for this, we performed parallel analyses on all neurons we recorded and also only on neurons with high SNR (> 3). We found the general pattern of results and our conclusions were consistent across these analyses, suggesting the results were robust to the particular criteria used for neuron type classification.

The continuum of time-varying attention effects we found in V4 is consistent with previous studies that have typically found variation in functional properties across neuron classes, but without clear categorical boundaries. For example, Hofer *et al*. (2011) found that the activity of excitatory and inhibitory ensembles depend on the their synaptic connectivity and selectivity to different visual features. Further, Kim *et al*. (2019) found that texture and shape tuning of V4 neurons lies on a continuum instead of strict boundaries between texture-preferring and shape-preferring neurons. Although it is natural to want to classify neurons or time-varying attention effects in explicit categories, this is likely an oversimplification of functional circuitry and may lead to misinterpreted results. Our work here associating this continuum with other physiological and functional properties of neurons imbues this variation in time-varying attention effects with renewed significance.

While it has long been appreciated that attention affects the firing rates of individual neurons (Luck *et al*., 1997; Moran & Desimone, 1985), there has been a growing appreciation of late for how correlated variability of neurons is affected in attention (Cohen & Maunsell, 2009; Mitchell *et al*., 2007; Smith & Kohn, 2008; Snyder *et al*., 2016; Snyder & Smith, 2018; Umakantha *et al*., 2021). For example, Cohen & Maunsell (2009) found that changes in interneuronal correlations better accounted for the behavioral effects of attention compared to firing rate correlates of attention. In another example, Mitchell *et al*. (2007) found that differential attention modulation of FS and RS neurons was not explained by differences in firing rates, but was better accounted for in terms of temporally-driven changes in interneuronal correlations. This latter result is especially relevant for the current report, where we also found differential attention effects across FS/RS neuron classes. In general, interneuronal correlations are important signatures of functional relationships between neurons, and they can be measured on slow time-scales (tens to hundreds of milliseconds, often termed “spike count correlations” in that case, as in Cohen & Maunsell (2009); Mitchell *et al*. (2009)) or on fast time-scales (< 10 milliseconds, termed “synchrony”), which we examined in the current study. We found that FS-FS neuron pairs are more synchronous than mixed (FS-RS) or RS-RS neuron pairs. This is in agreement with the proposed functional role of FS/putative inhibitory neuron synchrony in temporal coordination of responses (Cardin *et al*., 2009) and synaptic properties of inhibitory neurons (Bartos *et al*., 2007). One potential mechanism driving changes in neural synchrony is via modulation of local inhibitory networks by higher order brain regions such as PFC (Tiesinga *et al*., 2004; Pritchett *et al*., 2015; Vinck *et al*., 2013). These inhibitory networks contain a variety of FS neurons that can then act on RS neurons and other FS neurons to drive processing of visual stimuli and generate local field potential (LFP) oscillations (Agetsuma *et al*., 2018; Hasenstaub *et al*., 2005; Tiesinga *et al*., 2004; Shin & Moore, 2019; Hawken *et al*., 2020). For example, optogenetic activation of FS interneurons entrains gamma oscillations (Cardin *et al*., 2009; Pritchett *et al*., 2015), strengthening visual responses (Cardin *et al*., 2009) and enhancing perception of harder-to-detect sensory stimuli (an important behavioral benefit of attention) (Pritchett *et al*., 2015). Different attention modulation and synchronization of certain sub-populations can change sub-populations’ engagement in gamma oscillations and affect downstream sensory processing (Cardin *et al*., 2009; Fiebelkorn & Kastner, 2020). For example, spiking activity of RS and FS neurons are differently phase-locked to gamma-band LFPs (Vinck *et al*., 2013) and these neuron types differ in their sensitivity to feedforward and recurrent network connections (Hofer *et al*., 2011). Therefore, during attention, synchronization of local sub-populations of neurons in sensory cortical areas like V4 can drive gamma oscillations and amplify sensory information during stimulus processing (Fiebelkorn & Kastner, 2020; Shin & Moore, 2019).

While spike synchrony quantifies functional interactions between pairs of neurons, another important question is how an individual neuron is related to the activity of an entire population (“population coupling”), which may be difficult to infer directly from pair-wise metrics. Previous researchers examined how population coupling related to factors such as sensitivity to stimulus input or top-down modulation Okun. While acknowledging that population coupling spans a continuous spectrum, Okun *et al*. (2015) used the terms “soloists” and “choristers” to ease description of neurons at the low and high ends of that spectrum, respectively, as shall we. They found that choristers were more sensitive than soloists to a variety of inputs such as from sensory stimulation or top-down modulatory influence, and reasoned this enhanced sensitivity of choristers to input may reflect an amplification process of the local circuitry. The current findings add the neural correlates of attention to the context of that prior work. We found that neurons with “restless-weak” time-varying attention effects tended to have stronger population coupling than neurons with other patterns of time-varying attention effects. Our results suggest that choristers are relatively more active with attention (compared to with inattention) during spontaneous activity, but relatively less active with attention during stimulus responses. Although the magnitude of attention effects during spontaneous activity may be slight (on the order of a few spikes per second), a difference of even a single spike can have a substantial effect in the context of a coupled population (London *et al*., 2010). While further research will be needed to better understand the computational significance of these divergent effects of attention on choristers depending on stimulus context, one potential hypothesis is that local amplification during spontaneous activity may be beneficial for maintaining a cognitive state in the population (by amplifying top-down input), whereas local amplification during stimulus processing may be detrimental if bottom-up inputs are noisy.

There are several limitations of this study that will require future experiments to test hypotheses mentioned above. The behavioral task was designed with few repeats of each target stimulus and each trial happens on a very fast timescale. These design features enabled us to have very high statistical power to analyze responses to non-targets, but at the expense of lesser statistical power to analyze target stimuli. A future experiment with a greater proportion of target stimuli could help address the questions such as differences in processing target stimuli or the point at which the animal has accumulated enough sensory evidence to make a decision. We also used a small number of different stimulus orientations, again in order to have many repeats of identical stimuli. Thus, in this current study we do not have enough measurements in the orientation space to characterize changes in tuning curves of these neurons. Changes in tuning curves would add an additional important layer of information about time-varying attention effects. For example, previous research has shown that in visual cortex, FS neurons exhibit multiplicative gain in firing rates with attention (Nandy *et al*., 2017; Sani *et al*., 2017). Therefore, one role of FS neurons could be to control normalization that amplifies gain and/or sharpens tuning curves. Neither spontaneous activity in attention nor the role of neuron type has been adequately studied in non-human primates. While addressing these topics is necessary to understand attention circuitry, we still face technical limitations using this animal model. For example, we observed that certain FS neurons in our sample had both especially high synchrony and population coupling (sea-foam green sector in Figure 4b. We are unable to specifically isolate these neurons *in vivo* to determine if they are, for example, basket interneurons coupled by gap junctions, or are linked to gamma oscillations. Future work investigating how timevarying attention effectsof FS neurons relate to gamma oscillations in V4 and upstream cortical areas is necessary to link electrophysiology studies of attention in non-human primates to those investigating neural oscillations and human imaging studies.

In this study, we found a continuum of time-varying attention effects in V4 neurons. The pattern of attention dynamics depended on the neuron’s response during attention (suppressed or enhanced), neuronal identity (excitatory or inhibitory) and it’s functional connectivity. In particular, neurons that are restless-weak are more likely to be RS (putative excitatory) and have greater population coupling. These findings bring into focus a picture of local circuitry underlying the dynamic allocation of attention in which different computational roles are distributed across diverse mixtures of V4 neurons.

## Methods

### Ethical oversight

Experimental procedures were approved by the Institutional Animal Care and Use Committee of the University of Pittsburgh and were performed in accordance with the United States National Research Council’s *Guide for the Care and Use of Laboratory Animals*.

### Subjects

Two adult male rhesus macaques (*Macaca mulatta*) were used for this study. Surgeries were performed in aseptic conditions under isoflurane anesthesia. Opiate analgesics were used to minimize pain and discomfort perioperatively. A titanium head post was attached to the skull with titanium screws to immobilize the head during experiments. After each subject was trained to perform the spatial attention task, we implanted a 96-electrode Utah array (Blackrock Microsystems) in V4. The array was implanted in the right hemisphere V4 for Monkey P, and the left V4 for Monkey W. A detailed description of these methods and a separate analyses of a portion of these data were published previously (Snyder *et al*., 2018; Cowley *et al*., 2020; Snyder *et al*., 2021; Umakantha *et al*., 2021; Johnston *et al*., 2021).

### Array recordings

Signals from the arrays were band-pass filtered (0.3 - 7500 Hz), digitized at 30 kHz and amplified by a Grapevine system (Ripple). Signals crossing a threshold (periodically adjusted using a multiple of the root-mean-squared noise) were stored for offline analysis. We first performed a semi-supervised sorting procedure followed by manual refinement using custom MATLAB software (https://github.com/smithlabvision/spikesort), taking into account waveform shapes and interspike interval distributions. These initial sorting steps yielded 93.2 ± 8.9 (*mean ± SD*) candidate units per session in V4 for Monkey P and 61.9 ± 27.4 candidate units per session in V4 for Monkey W. We likely recorded a mixture of single units and multiunit activity, though for simplicity we refer to all units as “neurons”. The arrays were chronically implanted and likely recorded some (but not all) neurons over more than one recording session, but we calculated our results within each recording session and treated each session as an independent sample for the analysis.

### Receptive field (RF) mapping

Prior to beginning the behavioral task, we mapped the receptive fields (RFs) of the spiking neurons recorded on the V4 arrays by presenting small (*≈*1°) sinusoidal gratings (four orientations) at a grid of positions. We subsequently used Gabor stimuli scaled and positioned to roughly cover the aggregate RF area determined by the responses to the small gratings at the grid of positions. For Monkey P this was 7.02° full-width at half-maximum (FWHM) centered 7.02° below and 7.02° to the left of fixation, and for Monkey W this was 4.70° FWHM centered 2.35° below and 4.70° to the right of fixation. We next measured tuning curves by presenting gratings at the RF area with four orientations and a variety of spatial and temporal frequencies. For each subject we used full-contrast Gabor stimuli with a temporal and spatial frequency that evoked a robust response from the population overall (i.e., our stimulus was not optimized for any single neuron). For Monkey P this was 0.85 cycles/° and 8 cycles/s. For Monkey W this was 0.85 cycle/° and 7 cycles/s. For the task, we presented a Gabor stimulus at the estimated RF location, at the mirror-symmetric location in the opposite hemifield, or at both locations simultaneously.

### Behavioral task

Subjects maintained central fixation as sequences of Gabor stimuli were presented in one or both of the visual hemifields, and were rewarded with water or juice for detecting a change in orientation of one of the stimuli in the sequence (the target) and making a saccade to that stimulus (Figure 1). The probable target location was block-randomized such that 90% of the targets would occur in one hemifield until the subject made 80 correct detections in that block (including cue trials, described below), at which point the probable target location was changed to the opposite hemifield. The fixation point was a 0.6° yellow dot at the center of a flat-screen cathode ray tube monitor positioned 36 cm from the subjects’ eyes. The background of the display was 50% gray. We measured monitor luminance gamma functions by photometer and linearized the relationship between input voltage and output luminance using lookup tables. We tracked the gaze of the subjects using an infrared eye tracking system (EyeLink 1000; SR Research, Ottawa, Ontario). Gaze was monitored online by the experimental control software to ensure fixation within *∼*1°of the central fixation point throughout each trial. We excluded from analysis data segments during which a subject’s gaze left the fixation window. After fixating for a randomly chosen duration of 300 to 500 ms (uniformly distributed), a visual stimulus was presented for 400 ms, or until the subjects’ gaze left the fixation window, whichever came first. For the initial trials within a block, a Gabor stimulus was presented only in the hemifield that was chosen to have a high probability of target occurrence for the block. There were two cue conditions: a cue at the RF location (cue-RF) or a cue in the opposite hemifield (cue-away). These cue trials were to alert the subjects to a change in the probable target location and were excluded from the analysis. The initial cue location was counterbalanced across recording sessions. Once a subject correctly detected five orientation changes during the cue trials, bilateral Gabor stimuli were presented for the remainder of the block. Each trial consisted of a sequence of 400 ms stimulus presentations separated by 300–500 ms inter-stimulus intervals (uniformly distributed). Stimulus sequences continued until the subject made an eye movement (data during saccades were excluded from analysis), or a target was presented but the subject did not respond to it within 700 ms (i.e., a miss). For the first presentation in a sequence, the orientation of the stimulus at the cued location was randomly chosen to be 45° or 135° and the orientation of the stimulus in the opposite hemifield, if present, was orthogonal to this. Subsequent stimulus presentations in the sequence each had a fixed probability of containing a target (30% for monkey P, 40% for monkey W), i.e., a change in orientation of one of the Gabor stimuli compared to the preceding stimulus presentations in the trial. Within a block, 90% of targets (randomly chosen) occurred in one hemifield (valid targets) and 10% of targets occurred in the opposite hemifield (invalid targets). For valid targets, the orientation change was randomly chosen to be 1°, 3°, 6°, or 15° in either the clockwise or anticlockwise direction (monkey P: 11.49 ± 3.14 (mean ± SD, across sessions) valid targets of each orientation at each location; monkey W: 14.56 ± 4.75 valid targets of each orientation at each location). For invalid targets, the orientation change was always the near-threshold value of 3°, clockwise or anti-clockwise (because invalid targets occur infrequently, we restricted the number of orientation change magnitudes for this condition in order to derive a reasonable estimate of the target detection rate). We analyzed trials including either valid or invalid targets, but excluded from analysis all neural data from the time of target onset through the end of the trial. That is, we only analyzed responses to non-targets, of which there were two types: one with a 45° stimulus in the RF, and the other with a 135° stimulus in the RF. Monkey W completed 24 sessions of the experiment; monkey P completed 25 sessions. One session for each subject was subsequently excluded from analysis because of recording equipment failure.

### Single neuron responses

From the continuous recording, we extracted data segments from 300 ms prior to stimulus onset to 300 ms following stimulus offset (1 second total segment duration; Monkey P: 2253.1 ± 591.3 data segments per session, Monkey W: 2894.2 ± 946.8 data segments per session) and counted spikes for each neural unit in 1 ms bins. For the calculation of PSTHs (Figure 2), we smoothed spike trains with a 100 ms moving average prior to averaging across data segments. To test individual neurons for significant attention effects (i.e., a difference between the cue-RF and cue-away conditions), we binned counts for each data segment in 50 ms, non-overlapping time bins and compared cue-RF and cue-away conditions with a Wilcoxon rank-sum test. We tested for differences between cue conditions separately for each stimulus configuration (i.e., 45° or 135° stimulus in the RF). We considered a neuron to have a significant attention effect if at least two time bins were significant with *p* < 0.05 for either stimulus configuration. We excluded neurons without significant attention effects from all analyses except the analysis of synchrony (Figure 6). All recorded neurons were included for the synchrony analysis.

### Neuron-type Classification

Evidence suggests parvalbumin-expressing inhibitory interneurons typically have different action potential waveforms compared to other types of neurons, with relatively short latencies to peak depolarization (Barthó *et al*., 2004), brief hyperpolarization durations (Kawaguchi, 1993), and rapid peak rates of depolarization (McCormick *et al*., 1985). Neurons with these features are referred to as fast-spiking (FS) (sometimes also called “narrow spiking” in the literature) and classified as putative inhibitory neurons. This is in contrast to regular-spiking (RS) (sometimes also called “broad spiking”) neurons, classified as putative excitatory neurons. We caution that there are likely exceptions to these general trends, and ground truth of the neurotransmitter secreted by the neurons we recorded is currently unknowable for the data in this report. We classified neurons as FS or RS using maximum likelihood estimation of a mixture of Gaussians model based on multiple measures of waveform shape, a method that we described in detail previously (Snyder *et al*., 2016). In brief, for each neuron we approximated each average action potential waveform as a parametric function yielding estimates of the timing and width of negative and positive phases of the action potential, which we used to embed each neuron as a point in a three-dimensional feature space. The three features were (1) the time to the positive peak of the extracellular action potential, (2) the width of the positive phase of the extracellular action potential, and (3) the relative rate of the rising and falling phases of the initial voltage deflection of the action potential (Supplemental Figure 1). We then modeled the distribution of neurons in this feature space as a mixture of two Gaussian distributions, one representing the FS neurons and one representing the RS neurons. Maximally likely model parameters were fit using expectation maximization (using the ‘fit’ method for the MATLAB ‘gmdistribution’ class), using ten replicates of random initial conditions. The fitting algorithm converged on a consistent solution across the range of initial conditions tested. For fitting the model parameters, we used only the neurons with signal-to-noise ratios (SNR; ratio of the average waveform amplitude to the SD of the waveform noise) in the top quartile of our sample (*SNR* > 3.782), reasoning that these were most likely to be well-isolated single units. We then calculated the posterior probability that each neuron (regardless of SNR) came from either the FS or RS distribution. We considered neurons with a posterior probability greater than 0.5 of belonging to the FS distribution as an FS neuron. For all neurons (SNR > 1), 2354 were classified as FS and 1838 were significantly modulated with attention. 1302 neurons were classified as RS and 1043 of them had significant attention effects. For neurons with an SNR > 3, a total of 547 neurons were classified as FS, with 456 that had significant attention effects. 628 neurons were classified as RS, and 505 of these neurons had significant attention effects. We tested various thresholds around this classification boundary and found our qualitative pattern of results was highly consistent over substantial variation of this criterion.

### Quantifying diversity of attention effect time-courses

We limited our data set to neurons with at least two 50 ms time bins with significant effects of attention (see *Single neuron responses*). For each neuron, we calculated the time-course of attention effects by subtracting the average stimulus response during the cue-away condition from the average response of the cue-RF condition.

We first calculated time-varying attention effects separately for each stimulus configuration (45° or 135° stimulus in RF). Overall, time-varying attention effects were strongly correlated between the two stimulus conditions. For each neuron (SNR > 3) that had a significant attention effect in both stimulus conditions (see section *Single neuron responses*), we calculated the correlation over time of the time-varying attention effects between the two stimulus conditions. The average correlation across our sample of neurons was strongly positive (mean *r* = 0.21, *t*_2133_ = 32.92, *p* < 0.0001; Supplemental Figure 2). At the individual neuron level, 636 neurons (29%) had significant (*p* < 0.1 to be liberal) positive correlations of time-varying attention effects across stimulus conditions while only 44 neurons (2%) had significant (*p* > 0.1) negative correlations of time-varying attention effects across stimulus conditions (for 1490 neurons the correlation was not significant). Thus, it was reasonable to conclude that, in general, neurons’ time-varying attention effects were not dissimilar between stimulus conditions, and, in cases where neurons had significant attention effects for both stimulus configurations, we averaged attention modulation time courses across the two stimulus configurations. Otherwise, for neurons with significant attention differences for only one stimulus configuration, we considered only that stimulus configuration for subsequent analyses. If we restricted our entire analysis to only stimuli of one orientation or the other, the qualitative pattern of results and corresponding conclusions were unchanged (Supplemental Figure 2).

Then, to quantify the similarity in time-varying attention effects between pairs of neurons, we calculated Pearson distances (i.e., one minus the Pearson product moment correlation coefficient) between their attention effect time-courses. This metric achieves a minimum value of zero for a pair of neurons with perfectly correlated attention effect time-courses, and achieves a maximum of 2 for a pair of neurons with perfectly anti-correlated attention effect time-courses. We then reduced the dimensionality of our data using multidimensional scaling to two dimensions (i.e., embed the neurons in a two-dimensional space while reducing the distortion of their pairwise Pearson distances; MATLAB function ‘mdscale’; Figures 4a and 4b). In preliminary analyses we also tested embedding in other dimensions (three, four), and found our results and conclusions were highly similar to embedding in two dimensions. We first asked whether neurons were clustered in the resultant time-varying attention effects feature space. We tested this by using k-means clustering for models with various numbers of clusters and calculated the Bayes Information Criterion for the result (we also tested spectral clustering with k-means). None of the cluster models explained the data better than the model with a single cluster. Thus, we concluded that time-varying attention effects were not clustered but rather continuously variable. In the absence of intrinsic structure (clusters) for grouping neurons based on their time-varying attention effects, we arbitrarily grouped neurons by dividing the two-dimensional feature space into twelve sectors (Figure 4b). In preliminary analyses we tested a range of sector parcellation schemes (different sizes and offsets) and found the qualitative pattern of our results and conclusions were robust to moderate variation of the sectors used for grouping neurons; we determined that twelve sectors provided a good balance of dividing the space of attention effects finely enough to illustrate the range of the sample, while maintaining enough neurons within each group for adequate statistical power.

### Synchrony

For our analysis of synchrony (and population coupling), we included all V4 neurons, regardless of whether they showed significant attention effects. We computed cross-correlograms for all simultaneously-recorded neuron pairs, and defined synchrony as the integral of the cross-correlogram over lags from -10 ms to +10 ms To account for coincidental spiking that was due to chance, we computed jitter-corrected cross-correlograms (Kohn & Smith, 2005) with 20 ms jitter windows. Spikes were randomly shuffled among trials within each jitter window and cross-correlograms were recalculated. We subtracted the average synchrony from these spike-jittered cross-correlograms from our observed synchrony values, resulting in an estimate of excess coincident spikes per second beyond what would be expected by chance given the firing rates of the neurons (Figure 6). Because unusually high synchrony estimates would be parsimoniously explained by recording artifacts affecting multiple channels, we excluded outlying neuron pairs from subsequent analyses (synchrony greater than 4 standard deviations above the mean). We also excluded from synchrony analyses pairs of neurons recorded on the same channel.

### Population-coupling

We sought to quantify with a scalar value how synchronous each neuron was to the broader population. For this, some seemingly reasonable metrics, such as the average synchrony of a neuron to all other neurons in the sample, are in fact not ideal, as a neuron might have especially high synchrony with one or a small number of other neurons, leading to a high average synchrony, while remaining in a relatively isolated “clique” relative to the rest of the population. Instead, we favored the graph analytical metric of closeness centrality as our measure of population coupling, which takes into account the full structure of synchrony across the population. To this end, we represented our populations as graphs, with each neuron, *n*_*i*_, represented as a node, and the distance, *d*_*i,j*_, between nodes *n*_*i*_ and *n*_*j*_ weighted by the synchrony of the corresponding neuron pair (Figure 7b). Pairs of neurons with non-positive excess synchrony were not connected in the graph. Next, we reasoned that neuron pairs with higher synchrony are functionally “closer” than neuron pairs with lower synchrony. Thus, we specified the distance between nodes in the graph as the log-reciprocal of excess synchrony, *s*:

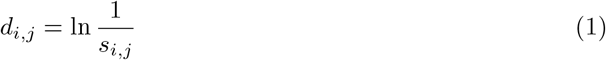

Note that *d* is not a *metric* in the formal mathematical sense, in that the triangle inequality is not necessarily satisfied. That is, the shortest path between a pair of neurons could be via a third neuron with which each neuron of the pair was more synchronous than they were with each other. To calculate closeness centrality of *n*_*i*_, we took the reciprocal of the average shortest path length of node *n*_*i*_ to all other nodes *n*_*j*≠*i*_ in the graph. Practically speaking, we used the ‘centrality’ method for graph objects in MATLAB.

### Statistical analyses

To test differences in behavioral accuracy (*d*′) and speed (RT) between validly-cued and invalidly-cued targets, we used two-tailed independent-samples Student’s t-tests with *α* = 0.05.

To test the responses of individual neurons for differences between cue-RF and cue-away conditions (i.e., attention effects), we used Wilcoxon rank-sum tests on spike counts in 50 ms non-overlapping bins with *p* < 0.05 in at least two time bins considered significant. Because we tested individual neurons for significant attention effects only for the purpose of inclusion/exclusion in subsequent analyses, we were intentionally liberal in our significance criterion and did not adjust for the family-wise error rate of testing multiple time bins (i.e., we wished to err on the side of including neurons, rather than excluding them).

To test for differences in odds ratio of FS neurons by MDS sector (Figure 5), we used a non-parametric bootstrap test. We first calculated the odds ratio that a neuron in each sector was FS relative to the overall risk that a neuron in the whole sample was FS:

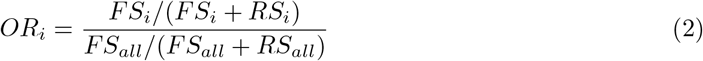

where *OR*_*i*_ is the odds ratio of an FS neuron in sector *i, FS*_*i*_ and *RS*_*i*_ are the numbers of FS and RS neurons, respectively, in sector *i*, and *FS*_*all*_ and *RS*_*all*_ are the numbers of FS and RS neurons overall, respectively.

We then compared this test statistic to a bootstrap distribution reflecting the null hypothesis that there was no difference in sector distribution between FS and RS neurons generated by randomly shuffling the FS/RS neuron-class labels and re-calculating *RR* for the label-shuffled distribution (repeated ten thousand times to create the distribution). The p-value for our bootstrap test was the proportion of *RR* values from the bootstrap reference distribution that was more extreme than the *RR* value observed for the original data. We used a type I error rate of *α* = 0.1, Sidak-corrected for the twelve of sectors tested (i.e., *p* < 0.0087), to determine significance.

To test for differences in synchrony as a function of neuron-type, cue condition, or stimulus context, we used a three-way omnibus ANOVA (Figure 6). We used factors of CLASS (three levels: FS-FS pairs, RS-rs pairs, and FS-RS mixed pairs), CUE (two levels: cue-RF/attended and cue-away/unattended), and STIMULUS (two levels: pre-stimulus and post-stimulus). We tested all main effects and interactions, followed by post-hoc pairwise comparisons with Tukey’s method, considering *p* < 0.05 as significant.

To test for differences in closeness centrality as a function of MDS (Figure 7c), we used a non-parametric bootstrap test. We first calculated the average closeness centrality for neurons in each sector as our test statistic (see section *Population-coupling*). We then compared this test statistic to a bootstrap distribution reflecting the null hypothesis that there was no relationship between closeness centrality and MDS sector generated by randomly shuffling the sector labels for the neurons and re-calculating the average closeness centrality in each sector for the label-shuffled data (repeated one thousand times to create the distribution). We considered observed values more extreme than 95% of the values in the bootstrap reference distribution to be significant.

## Acknowledgements

A.C.S. was supported by NIH grants K99/R00EY025768, R01EY028811 and R01EY011749; a NARSAD Young Investigator award from the Brain & Behavior Research Foundation; and an Alfred P. Sloan Foundation research fellowship. Experimentation was performed by A.C.S. while a postdoctoral fellow in the laboratory of Matthew A. Smith at the University of Pittsburgh (now at Carnegie Mellon University). We thank Dr. Smith for his support, including funding (NIH grants R01MH118929, R01EB026953, R01EY022928 and P30EY008098; NSF NCS BCS 1954107/1734916; Research to Prevent Blindness; and the Eye and Ear Foundation of Pittsburgh). The authors would like to thank Ms. Samantha Schmitt for assistance with surgery and data collection.

