## Supplemental Figures for "Dynamic attention signaling in V4: relation to excitatory/inhibitory cell class and population coupling"

distribution of waveform classification parameters, SNR > 3.0

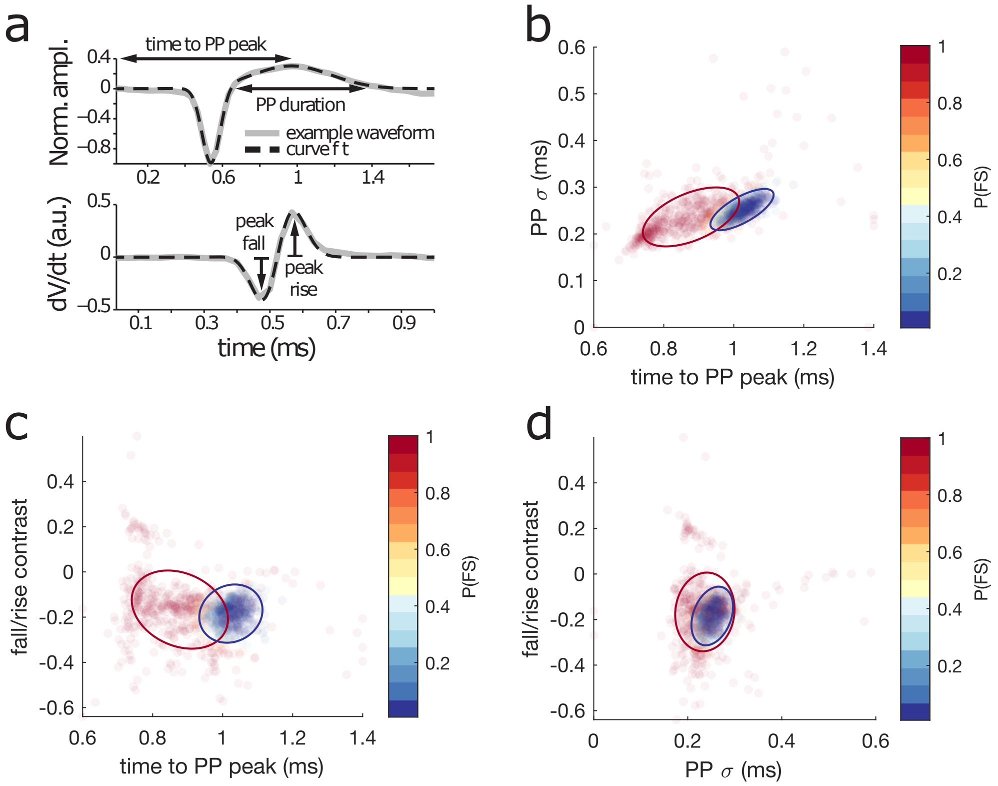


Supplemental Figure 1: distribution of waveform shape parameters for V4 neurons with signal-to-noise (SNR) greater than 3, and results of classification procedure. a) shape parameters. To quantify waveform shapes, we approximated each waveform by fitting to a sum of two Gaussian functions (cf. Snyder et al., 2016 for detailed methods). We used the shift and standard deviation of the positive Gaussian as the (1) time to the positive phase (PP) peak, and (2) duration of the PP, respectively. We used the difference in amplitude of the positive and negative peaks of the waveform derivative as the fall/rise contrast. These three waveform metrics have previously been shown to differ between fast-spiking and regular-spiking neurons (Kawaguchi, 1993; McCormick et al., 1985). b-d) two-dimensional marginal distributions for our three-dimensional waveform data. Each dot represents a neuron, and the color of each dot represents that neuron's posterior probability of belonging to the fast-spiking class. Ellipses enclose confidence intervals for the mean of each class distribution (red: fast-spiking, 50% CI, blue: regular-spiking, 95% CI). We classified neurons using a Gaussian mixture model, assuming two classes, where the Gaussian component parameters were estimated by fitting to the sample of neurons in the top quartile of SNR (Snyder et al., 2016).

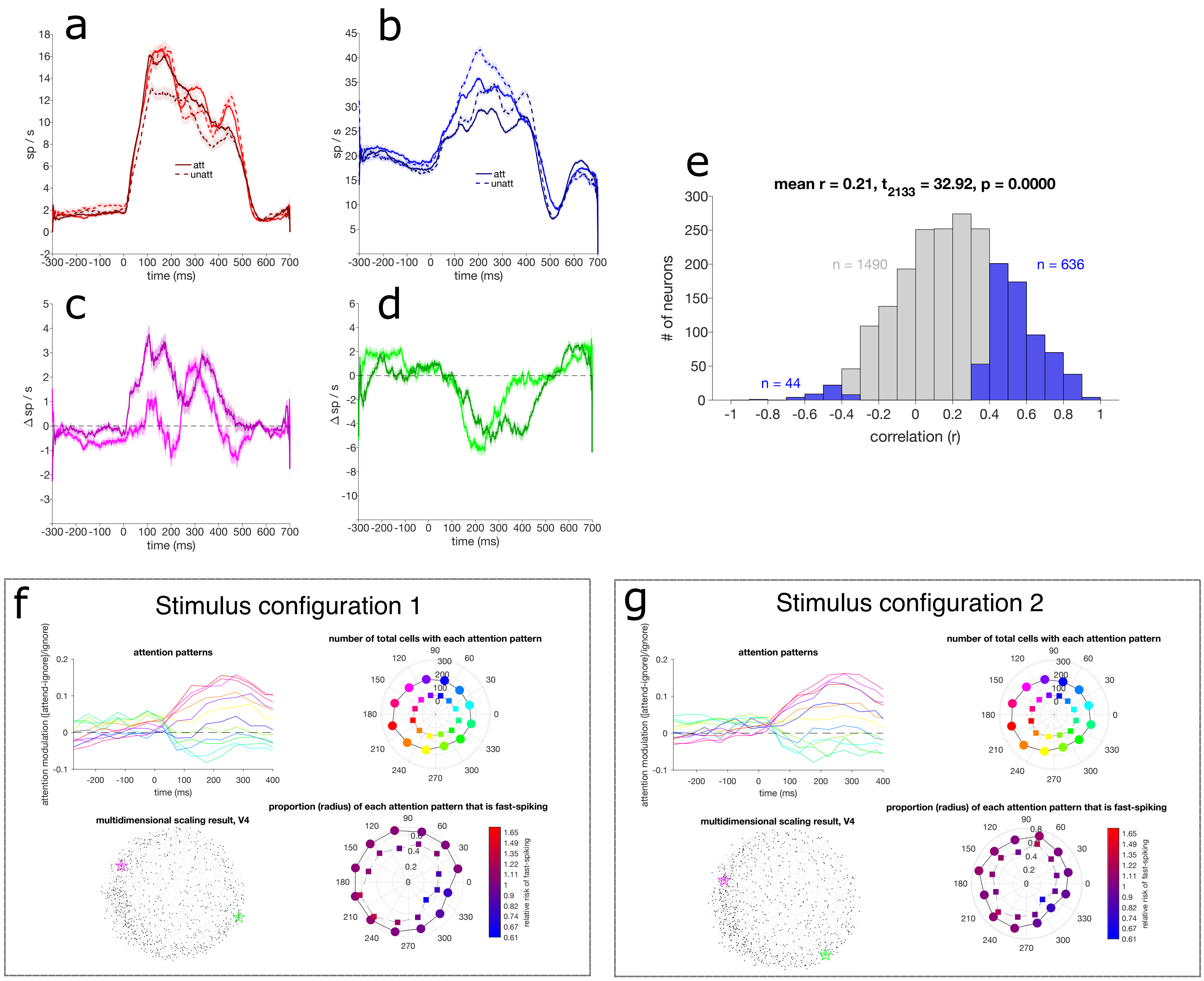


Supplemental Figure 2: the pattern of results and our conclusions are similar if analyses are restricted to just one stimulus configuration or another. For "stimulus configuration 1", the 45 degree grating was in the receptive field, while for "stimulus configuration 2" the 135 degree grating was in the receptive field. a-b) Example neurons' (same as in Figures 2 & 4 of main report) PSTHs for attended (solid) and unattended (dashed) conditions for stimulus configuration 1 (lighter colors) and configuration 2 (darker colors). c-d) Difference between attention conditions for same neurons as in 'a' and 'b', respectively, for stimulus configuration 1 (lighter colors) and configuration 2 (darker colors). Note general similarity of time-course (e.g., negative before stimulus onset at time 0s, and generally positive after stimulus onset in panel 'c', vice versa for panel d). e) Across neurons, attention effect time-courses are generally positively correlated between stimulus configurations. For each neuron that had a significant attention effect for each stimulus configuration (see methods for determination of significant attention effects), we calculated the Pearson correlation in attention effect time courses between stimulus configurations. The overall distribution of correlations was significantly positive (r = 0.21, p<<0.001). For individual neurons with significant correlation in attention effect time-courses between stimulus configurations, almost all of them were positively correlated (636 out of 680 neurons). Taken together, this pattern of results indicates that, at least, attention effect time-courses are not generally different between stimulus configurations. f-g) Multidimensional scaling of attention effect time-course distribution results are similar when restricted to one stimulus configuration ('f') or another ('g'). Same analyses as in Figures 4 & 5 of the main report.
